# Analysing the role of SERPINE1 network in the pathogenesis of human glioblastoma

**DOI:** 10.1101/2022.12.19.520990

**Authors:** Zahra Khosravi, Chandrasekaran Kaliaperumal, Arun HS Kumar

## Abstract

**Background:** Glioblastoma multiforme (GBM) is one of the most aggressive and difficult to treat brain tumour in humans with a 5 year survival rate of less than 6%. SERPINE1 is a novel tumour receptor found on GBM that modulates the progression of this cancer through growth signals and remodelling of the extracellular matrix. Hence, we investigated the role of SERPINE1 and its network proteins in pathogenesis of GBM and assessed its targetability.

**Material and methods:** Network proteins of SERPINE1 in homo sapiens was identified using the String database, and the affinity of the protein-protein interaction of this network was analysed using Chimera software. The expression profile of SERPINE1 in the different brain regions was evaluated to correlate its relevance to GBM pathology. Selected small molecules from *Calotropis gigantea* were screened using AutoDock vina to assess targetability of human SERPINE1.

**Results:** VTN, PLG, TGFB1, VWF, FGF2 and CXCR1 were identified as the major network proteins of SERPINE1. The strongest interaction was observed between SERPINE1 and FGF2 (42884 H-bonds) followed by CXCR1 (20279 H-bonds). Our results suggest that SERPINE1 and its network proteins identified here play a vital role in GBM development and progression through brain parenchyma by creating the prime microenvironment for carcinogenesis, tumour invasion and migration. The highest expression of SERPINE1 was observed in the pons, medulla, midbrain, corpus callosum and spinal cord. Expression of SERPINE1 was consistent with high grade lesions of GBM, suggesting association of SERPINE1 with advanced stages of GBM. The selected small molecules from *Calotropis gigantea* were observed to have therapeutically feasible binding affinity (140 - 550 μM) and predicted efficacy (290 - 1115 μM) against human SERPINE1.

**Conclusion:** SERPINE1 plays a vital role in the progression of GBM through its critical network proteins identified in this study. The expression of SERPINE1 aligns with the advanced stages of GBM. Small molecules from *Calotropis gigantea* tested in this study can serve as lead compounds for developing novel anti-SERPINE1 therapeutics for advanced stages of GBM.

## Introduction

Glioblastoma multiforme (GBM) is an aggressive type IV cancer^[1, 2]^ with median survival duration of 15-23 months and less than 6% 5-year survival rate.^[2, 3]^ The tumour arises from astrocytes but significantly affects the glial cells surrounding neurons.^[1–3]^ Contributing to the hostile nature of this tumour are finger-like projections that heterogeneously extend into surrounding brain areas which make this tumour a challenge to treat on account of its poor localization and difficult anatomical access.^[1–31]^ The poor localisation of GBM further contributes to the difficulties in performing post-surgical resection and hence its higher incidence of recurrence. GBM exhibits considerable intra and inter-tumour heterogeneity, with as much as 50% of recurrent tumour samples showing diverse genetic mutations from the primary tumour.^[4, 5]^ Gliomas can also potentially differentiate from myeloid progenitor cells, which are prevalent within the tumour microenvironment^[6]^ and contribute to the tumour upregulation in diffused neurogenic zones. The genetic heterogeneity and cellular diversity of GBM adds to the therapeutic and surgical challenges faced. Additionally, resistance to chemotherapy, radiation and immunotherapy as an ongoing process through increasing expression of efflux transporters and immune suppression mechanisms account for poor clinical outcomes.^[4, 5]^ The invasive nature of GBM is strongly stimulated by cytokines which are an essential regulators of epithelial-mesenchymal transition (EMT).^[1]^ Among EMT program genes which were upregulated, SERPINE1 was also markedly elevated in the dispersive, motile population of GBM cells.^[1]^ These cells are prevalent in the mesenchymal subtype of GBM, which also involve NF-kB pathways.^[1, 4]^ This subtype of GBM is considered as a resistant phenotype which corresponds to poor survival and enhanced invasiveness.^[1]^

SERPINE1 is a regulator of the plasminogen activator system, with a significant role in the remodelling and degradation of the extracellular matrix.^[1, 4]^ Its importance in GBM progression is due to fibrinolysis inhibition. TGFß signalling controls the expression of SERPINE1, which inhibits fibrinolysis leading to migration of GBM cells, cell-adhesion and angiogenesis.^[1, 4]^ As an adipokine, SERPINE1 acts in an autocrine, paracrine and endocrine manner and leads to cancer progression through proinflammatory and anti-apoptotic pathways.^[7]^ SERPINE1 is believed to be a primary inhibitor for urokinase-type plasminogen activator (uPA) which ultimately leads to cell migration and proliferation.^[8]^ Thus, the mesenchymal subtype of GBM is enriched with SERPINE1.^[1, 2]^ Its significance in GBM progression was evident from how the knock-down of SERPINE1 reduced dispersal and focal adhesions to the extracellular environment.^[1, 2]^ Silencing of SERPINE1 was reported to reduce the expression of VEGFA in triple-negative breast cancer cells.^[9]^ This effect, in addition to Fas/FasL-dependent apoptosis suppression involves the EGFR, Akt and ERK signalling pathways. Additionally, SERPINE1 is also reported to be involved in development of resistance to paclitaxel.^[9]^ These attributes indicate that SERPINE1 could be a potential drug target against solid tumours including GBM. SK-216, which is a specific SERPINE1 inhibitor, was reported to reduce tumour progression in mice with Lewis lung carcinoma and malignant pleural mesothelioma.^[8, 10]^ Thus, it can be inferred that SERPINE1 has an important role in tumour progression which is why the focus of this study was to analyse the network proteins of SERPINE1 in order to exemplify its relevance to pathophysiology and targetability of GBM.

## Materials and Methods

A network protein analysis of SERPINE1 in Homo Sapiens was conducted using the STRING Database (https://string-db.org), to observe its functional protein-protein interactions.^[11–14]^ The protein networks were identified and reviewed in the UniProt database and the most resolved protein structure based on the full length of protein and atomic resolution was selected for analysis in this study. The PDB code of the selected protein structure was used to downloaded the structure onto the Chimera software and the number of hydrogen bonds (H-bond) formed between SERPINE1 and its associated network proteins at 2, 5 and 10 Armstrong distance was evaluated. A heatmap of the number of H-bonds formed between SERPINE1 and its associated network proteins was generated to identify the high affinity interactions. The expression profile of SERPINE1 in different areas of brain tissue were text mined from the Human Protein Atlas (https://www.proteinatlas.org) database and analysed by correlating the expression profile with severity of GBM. As the protein expression data of SERPINE1 in various brain regions is lacking we looked at the normalised expression of mRNA data with an assumption that this will reflect the protein expression pattern of SERPINE1 in different regions of the human brain.

To assess the targetability of SERPINE1 by small molecules, in this study molecular docking of selected molecules from *Calotropis gigantea* (Nicotiflorin, mefruside, quercetin, and zingerone) against crystal structure of human specific SERPINE1 was performed using AutoDock vina 1.2.0. Briefly, the PDB structure of SERPINE1 was optimised for molecular docking in the AutoDock MGL tools. The isomeric SMILES sequence of the selected molecules from *Calotropis gigantea* were obtained from the PubChem database and were converted into the PDB format using the Chimera software.^[15]^ The PDB structures of *Calotropis gigantea* molecules were further processed in the AutoDock MGL tools for molecular docking as reported previously,^[11, 13, 14]^ Imatinib was used as a reference compound.

## Results

The primary network analysis of SERPINE1 is summarised in figure 1. Besides these 10 network proteins of SERPINE1 a few more myeloid cell specific targets were considered for protein-protein hydrogen bond (H-bond) interaction analysis in this study. The H-bond analysis carried out is summarised in figure 1. It can be observed that VTN, PLG, TGFB1, VWF, FGF2 and CXCR1 had the highest interaction with SERPINE1. The strongest interaction was observed between SERPINE1 and FGF2 (42884 H-bonds) followed by CXCR1 (20279 H-bonds). These proteins are involved in regulation of inflammation and angiogenesis respectively and hence may be relevant to the development and progression of GBM through brain parenchyma. As a key mitogenic factor in tissue homeostasis, FGF is involved in the regulation of a variety of stem cells through receptor tyrosine kinases. Specifically, it has been shown that it is required in early neural development and neural stem cell expansion.^[16, 17]^ A positive correlation between glioma progression and FGF1 and 2 expression is also reported. Although FGF2 signalling in GBM is not yet fully understood, FGF1 is known to promote tumorigenicity and FGF2-specific anti-sense oligonucleotides are reported to halt glioma cell proliferation and angiogenesis in vitro.^[16, 17]^ The RAS signal-regulated pathways are also hyperactivated in GBM through the binding of FGF2 to FGFR1 on cancer stem cells. Therefore, FGF ligand/receptor signalling pathways play an important role in promoting GBM survival and expansion and perhaps this is influenced upstream by SERPINE1.

**Figure 1:**
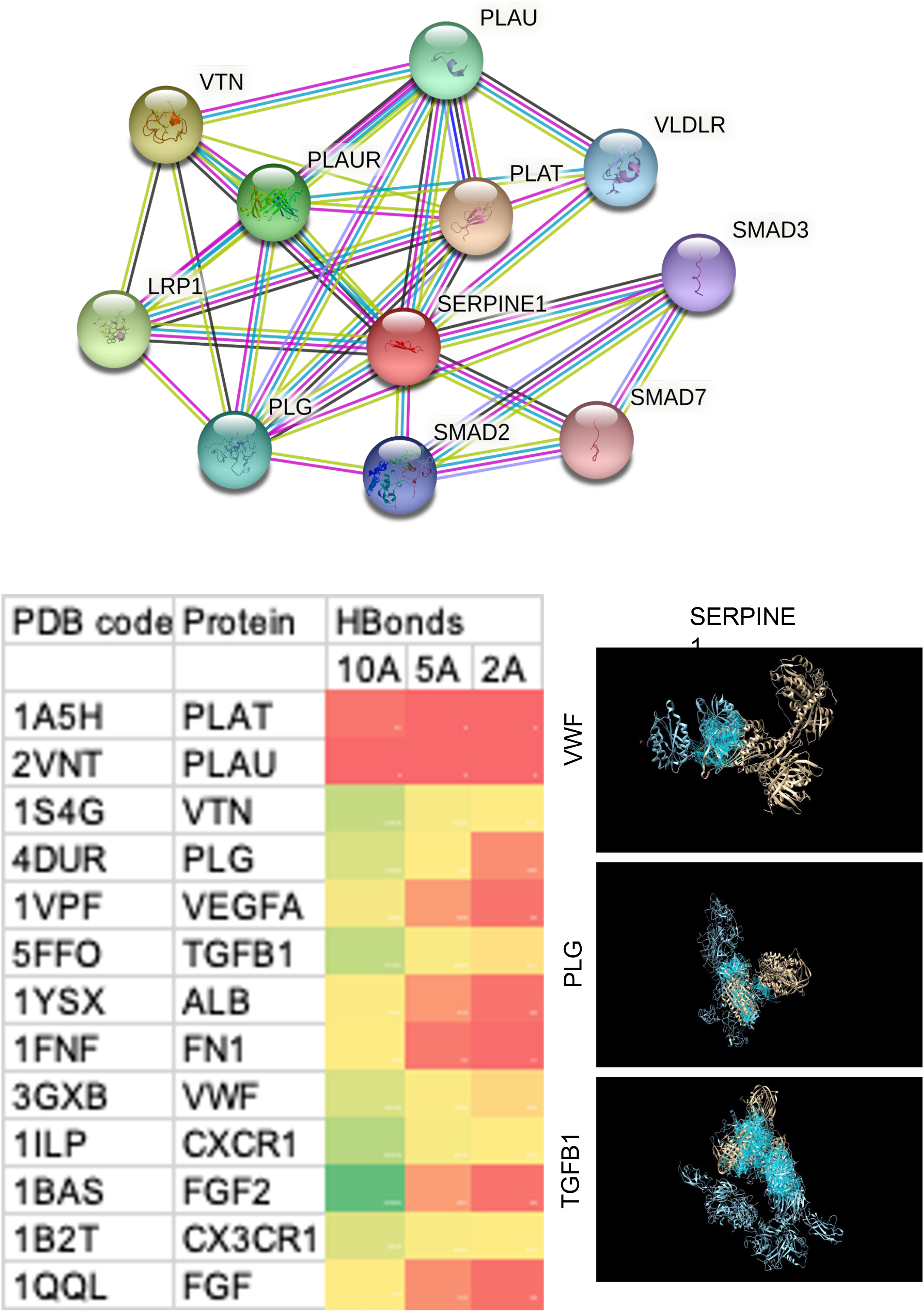
The major network proteins of SERPINE1 in humans. Heatmap showing the degree of interaction of each of the network protein with SERPINE1 at variable hydrogen bond (HBond) distance (2, 5 and 10 Armstrong). Selected human specific network proteins (TGFbeta1, VWF and PLG) in cyan are show interacting (blue lines represent the hydrogen bonds) with human SERPINE1 (brown).

CXCR1 and CX3CR1 are myeloid specific receptors activated by IL-8 and are known angiogenic factors. SERPINE1 was observed to have significant affinity to both these factors in this study. IL-8 is upregulated in GBM and its association with CXCR1 is observed in neoplastic vessels.^[18]^ IL-8 is also a downstream activator of STAT3, which ultimately maintains the stem cell potency of GBM cells and promotes GBM angiogenesis and invasion.^[19]^ Inhibition of CXCR1 by neutralising antibody showed significant decrease in GBM neurosphere formation and colony forming ability.^[19]^ Supporting the pivotal role of CXCR1 mediated STAT3 signalling in GBM stem cells, is the report of cancer stem cell depletion in human breast cancer by blocking CXCR1.^[20]^ The higher number of H bonds observed between SERPINE1 and CXCR1 seems to be significant in GBM as SERPINE1 being upstream to CXCR1 may be promoting pro-tumorigenic and pro-angiogenic pathways necessary for expansion of GBM. TGFß signalling is dysregulated in cancer signalling cascades. Although in normal epithelial cells it acts as a growth inhibitor, GBM cells show resistance to TGFß induced growth inhibition due to SMA, PI3K and FoxG1 alterations.^[21]^ TGFß also promotes invasion and migration of GBM cells through recruitment of macroglia (which express CXCR1 and CX3CR1) that constitute most of the tumour-infiltrating immune cells in GBM.^[22]^ TGFß also mediates immunosuppression by inhibiting the expressions of perforin, granzyme A and B and Fas ligand which are all responsible for cytotoxic T-lymphocyte mediated tumour cytotoxicity.^[21, 22]^ Later in the carcinogenesis process, TGFß is secreted in high amounts by the tumour stromal cells, creating a favourable environment for angiogenesis.^[21, 22]^ Therefore, the observation of significant interaction between SERPINE1 and TGFß1/ CXCR1/CX3CR1 in this study reflects the role of this network in the invasiveness and perhaps aggressiveness of GBM.

With 16212 H-bonds with SERPINE, vitronectin (VTN) was another protein of significance in the SERPINE1 network. VTN is an extracellular matrix protein that has a role in determining glioma tumour grade.^[22]^ Furthermore, its permissive role as a substrate for glioma migration has been established^[23]^ within the CSF and serum. The antiapoptotic effects of VTN through inhibition of topoisomerase and increasing Bcl-2 and Bcl-x proteins at the tumour-brain interface confers a survival advantage and invasiveness to GBM cells.^[24]^ On that account, SERPINE1 and its high affinity network proteins seem to validate why GBM has a high rate of invasiveness and tumour burden. SERPINE1 was also observed to have high affinity with PLG and VWF, both of which are essential regulators of haemostasis and have vital role in inflammation, inflammatory cell migration, interaction with complement proteins and extracellular matrix degradation.^[1, 25]^ PLG is reported to be involved in the growth and angiogenesis in malignant form of gliomas^[26]^ perhaps by facilitating extracellular matrix invasion. Hence PLG together with VWF and VTN can promote the SERPINE1 regulated GBM invasion.

In addition to analysing the network proteins of SERPINE1, this study also evaluated the expression profile of SERPINE1 in different regions of the human brain (figure 2). The highest expression of SERPINE1 was observed in the pons, medulla, midbrain and corpus callosum. Incidentally significant expression of SERPINE1 was also observed in the spinal cord. The expression pattern of SERPINE1 didn’t fully align with the general localisation of GBM, which is reported to be most commonly located in the frontal and temporal lobes.^[27]^ However high grade lesions of GBM are known to be localised in pons, midbrain and spinal cord, which perhaps reflects the association of SERPINE1 with advanced stages of GBM. Furthermore the biochemical role of SERPINE1 in haemostasis, migration and remodelling is consistent with SERPINE1 supporting the advancement of GBM progression and severity. The expression pattern of SERPINE1 observed in this study perhaps reflects its role in the invasion and spread of GBM to critical brain regions that regulate vital autonomic functions and hence the disease severity with associated high mortality. The observation in this study associating SERPINE1 with advanced GBM lesions warrants examining approaches of targeting SERPINE1 in clinical management of advanced stages of GBM. Epithelial to mesenchymal transition (EMT) marks the increase in the invasiveness and migration of cancer cells.^[1, 27]^ The role of SERPINE1 in EMT and tumour invasiveness/aggressiveness is well known^[27]^ in several types of cancers including triple negative breast cancer and gastric cancer.^[27]^ Such direct associations further support the relevance of therapeutically targeting SERPINE1 in clinical management of advanced stages of GBM.

**Figure 2:**
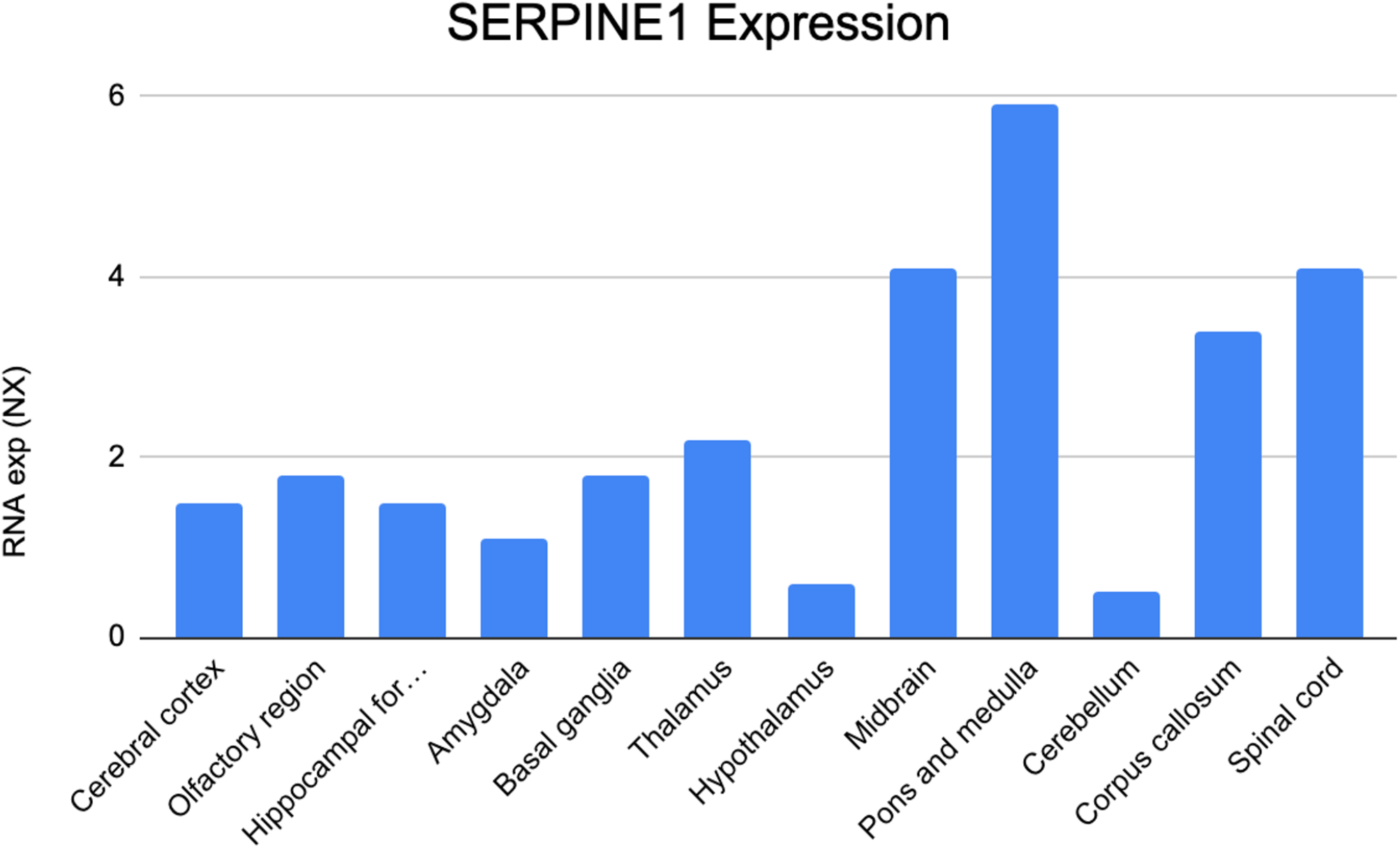
The bar graphs represent normalised expression (NX) of SERPINE1 mRNA in various regions of the brain.

We have recently reported the utility of phytochemicals from *Calotropis gigantea* as potential anticancer therapeutic leads.^[15]^ Here we investigated selected compounds from *Calotropis gigantea* for their feasibility to target SERPINE1. The binding affinity (140 - 550 μM) and predicted efficacy (290 - 1115 μM) of the compounds with human SERPINE1 were observed to be in the therapeutic range (figure 3). Nicotiflorin and quercetin showed highest efficacy, followed by zingerone and mefruside. All the four phytochemicals from *C.Gigantea* showed higher binding affinities to SERPINE1 than Imatinib. Several clinical trials have examined the potential of Imatinib for the treatment of GBM, with limited efficacy.^[28–30]^ Considering the superior efficacy of *C.Gigantea* compounds to target SERPINE1, these compounds can be further structurally refined and improved for clinical use.

**Figure 3:**
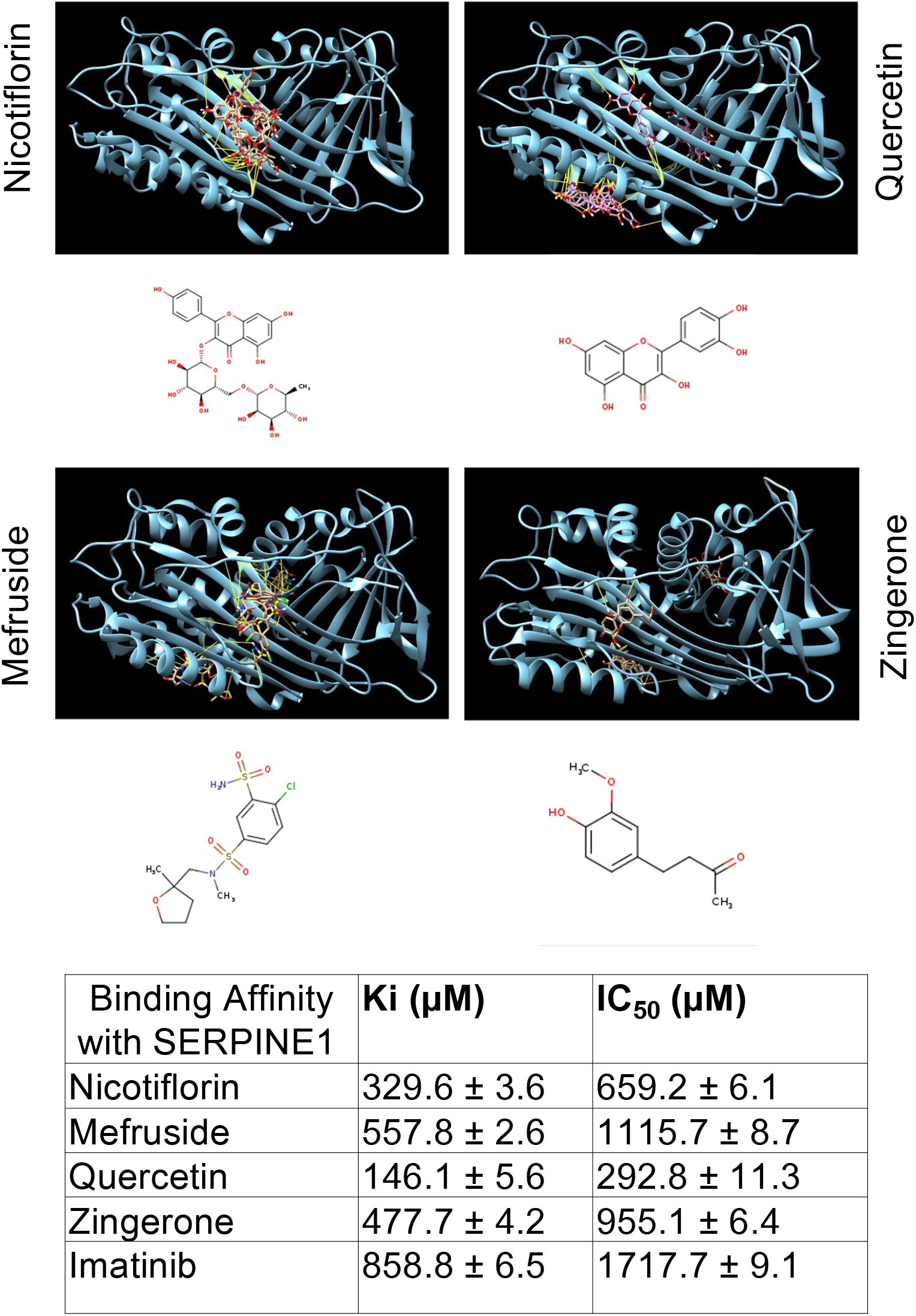
Molecular docking of selected small molecules from *Calotropis gigantea* (Nicotiflorin, mefruside, quercetin, and zingerone) against human SERPINE1 (Cyan). Images show the site of interaction (yellow line represent hydrogen bonds) between the small molecule and human SERPINE1. The binding affinity (Ki) and efficacy (IC_50_) of these small molecules to inhibit SERPINE1 is represented in the table.

## Discussion

The sinister nature of GBM stems from its ability to locally invade and disperse within brain tissue. There are several growth factors and genes that give GBM its plasticity and of these, SERPINE1 through its network proteins plays an important role as highlighted in this study. Specifically the network protein analysis of SERPINE1 showed its molecular interactions with key proteins reported to be involved in aggressiveness of GBM. The role of SERPINE1 with advanced forms of GBM was further evident from its expression analysis in brain tissue, which revealed higher association with brain regions involved in regulation of vital autonomic physiology. This observation is consistent with clinical reports suggesting significant association of SERPINE1, NAMPT, GRN, IL6 and CCL2 expressions with higher grade GBM.^[7]^ In our network protein analysis SERPINE1 was observed to be significantly associated with FGF2, CXCR1 and CX3CR1, all of which similar to CCL2 represent myeloid cell morphology. These associations in our view promote tumour invasiveness and aggressiveness by supporting EMT, angiogenesis and tumour cell plasticity. Consistent with this conclusion, knockdown of SERPINE1 was shown to suppress tumour invasion, growth and dispersion from the primary tumour core.^[7, 31]^ In contrast overexpression of SERPINE1 was found to be more prevalent in grade III gliomas,^[7, 31]^ drawing in the relevancy of our results of SERPINE1 expression in the critical regions of brain reflecting advanced stages of GBM.

SERPINE1 expression in gastric carcinoma, was reported to upregulate VEGF and IL-6 which lead to increased migration and invasion of cancer cells.^[32]^ Consistent with this higher SERPINE1 expression was also found specifically in invasive forms of GBM.^[33]^ High grade brainstem gliomas compromise only 1-2% of GBMs, but carry a very high burden due to their proximity to delicate structures.^[34]^ Due to this, surgical resection is often deemed challenging, which is why investigating different methods of therapy is necessary. Based on the observation from this study, one such alternative approach could be by selectively targeting SERPINE1 using small molecules. GBM displays increased cell density, cellular atypia and a necrotic core which are supported by angiogenesis promoted by VEGF upregulation.^[4]^ Unlike VEGF which is involved in physiological and pathological angiogenesis, targeting of SERPINE1 interferes with pathological angiogenesis and hence it offers a selective and specific therapeutic approach. Further vasculogenesis through involvement of myeloid cells disrupts the normal physiological barrier, causing blood vessel leakiness and potential collateral pathophysiology.^[3]^ Targeting of SERPINE1 is likely to impact several such collateral pathophysiology associated with severity of GBM. Tumorigenesis and its evasion can also be influenced by cellular energetics^[5]^ and targeting of SERPINE1 can interfere with cellular energetics of GBM and improve therapeutic efficacy.

Currently temozolomide is a first-line treatment option for GBM, however almost all GBM responding to first line therapy recur.^[4, 5]^ Most GBM exhibit EGFR amplification, however EGFR inhibitors have failed in clinical trials due to intratumoral heterogeneity and poor blood-brain-barrier (BBB) permeability.^[4, 5]^ Furthermore the modulation of BBB by glioma stem cell transcription factors confer to GBM chemoresistance^[4–6]^ Currently there is a significant lack of treatment options for GBM and several novel therapeutics approaches^[4, 28]^ including phytochemicals^[35]^ are being explored. Hence to address this therapeutic gap, having established the merit of targeting SERPINE1 for the treatment of GBM, in this study we propose potential lead-like molecules derived from *Calotropis gigantea* which can be developed for clinical use. Four compounds from *C gigantea* showed high efficacy against human SERPINE1 within therapeutically feasible concentration, which in our opinion can be further developed for clinical use.

## Acknowledgement

Research support from University College Dublin-Seed funding/Output Based Research Support Scheme (R19862, 2019), Royal Society-UK (IES\R2\181067, 2018) and Stemcology (STGY2917, 2022) is acknowledged.

